# GPU-accelerated, self-optimizing processing for 3D multiplexed iterative RNA-FISH experiments

**DOI:** 10.1101/2025.10.10.681751

**Authors:** Rory Kruithoff, Mauri D. Spendlove, Steven J. Sheppard, Maxwell C. Schweiger, Pedro Pessoa, Mohammad Abbasi, Steve Pressé, Benjamin B. Bartelle, Douglas P. Shepherd

## Abstract

Imaging-based spatial transcriptomic approaches rely on iterative labeling and imaging of carefully prepared samples, followed by solving a computational inverse problem to determine the location and identity of the targeted RNA. Because these approaches require high-resolution optics, the Nyquist-Shannon determined voxel size is small relative to typical tissue sample footprints. A common solution to speed up both experiments and computation is to increase the distance in between focal planes, trading off local information content to sample a larger imaging area in a reasonable time. In this work we introduce a GPU-accelerated computational framework, `merfish3d-analysis`, designed to speed up the computational processing of barcoded, in situ imaging-based spatial transcriptomics. Using this framework, we quantify the information lost due to axial sampling changes in simulated imaging-based spatial transcriptomic experiments, robustly reprocess publicly available multiplexed error-robust fluorescence in situ hybridization (MERFISH) datasets, and analyze new MERFISH experiments performed on a post-mortem human olfactory bulb sample. To improve the quality of experimental data in the post-mortem human sample, we designed a multi-step autofluorescence quenching protocol specific for in situ imaging-based spatial transcriptomic strategies. Taken together, we hope that the sample preparation protocols and single workstation, GPU-accelerated processing will further democratize imaging-based spatial transcriptomic experiments.

## INTRODUCTION

Gene expression dynamics involves the 3D progression of molecular signaling and structural changes as cell types emerge and self-organize into functional units. High-throughput and high-content molecular methods, such as DNA sequencing [1], single-cell RNA sequencing (scRNA-seq [2]), and cytometry by time-of-flight (CyTOF [3]), have transformed our understanding of the molecular gene expression underlying single-cell identity [4, 5]. Computational methods allow for estimation of cell identity, time correlations, and trajectories from these dissociated collections of individual cells [6]. Because such standard “-omics” methods lack spatial information, further technologies have focused on performing high-content measurements in situ, with varying levels of molecular content, or “plexity” and spatial resolution [5, 7, 8].

In the spatial “-omics” field, various approaches exist to multiplex from tens to thousands of gene targets using imaging-based transcriptomics [9–13] or whole transcriptomes using sequencing-based approaches [14–16]. For imaging-based approaches that rely on sequential single-molecule labeling, detection, and decoding by iterative label exchange, such as multiplexed error-robust fluorescence in situ hybridization (MERFISH [10, 17, 18]), high-resolution optical microscopes are paired with computational approaches to resolve individually labeled molecules, as well as compatibility with fluidic cells to perform the required chemistry for label exchange.

Recent efforts have expanded the 2D field of view (FOV), axial depth, axial sampling, number of measurable species (“plexity”), and signal amplification through a combination of biochemical, computational, and optical improvements [19–21]. Alternatively, the “plexity” can be increased through computational methods to impute, or infer, unmeasured genes from reference scRNA-seq atlases [22–26]. Significant challenges remain in applying these innovations to large samples of aged human tissue, which are often burdened by overwhelming autofluorescence and can lead to inaccurate decoding [27].

Ultimately, computational decoding of imaging-based spatial transcriptomics experiments is an inverse problem with the goal of estimating the transcripts present in the sample given the data, prior information about the experimental design, and physics of the imaging system. There are multiple strategies to perform this inference, ranging from simple estimators to more sophisticated Bayesian schemes. A non-exhaustive list of prior work in this area includes highperformance 2D pixel-decoding [28–30], 2D/3D spot finding and linking [13, 31], maximum likelihood approaches [32, 33], 2D deep learning [34], and optimization-driven approaches [35– 37]. There are also many approaches to finding and localizing diffraction-limited features, thanks to the single-molecule localization community [38].

Most of these approaches treat each axial plane in the acquired datasets as independent from other planes. This assumption is usually justified, as the axial spacing z’ of the acquired data is significantly larger than the Nyquist-Shannon sampling limit for the given microscope configuration, for example z’=1.5 µm instead of z’*∼*0.3 µm for a high numerical aperture (NA) oil objective. Acquiring and processing properly sampled 3D data increases experimental and computational costs, as the imaging time extends and the image processing, decoding, and filtering steps now include a third dimension. The increase in time cost leads to a trade-off in the number of cells imaged vs the density of the measurement within each cell. The trade-off is evident in large-scale atlasing efforts, where very few axial planes with large interplane distance are measured [39, 40] or samples were effectively “flattened” into one axial plane by digestion before imaging [41].

Beyond the sampling strategy, the sample quality will also impact the number and spatial distribution of recovered RNA. For ideal tissue types with minimal autofluorescence, sample preparation methods incorporating perfusion fixation to enhance RNA preservation have been shown to yield RNA counts in the hundreds per cell. While no absolute ground truth measurement is available, these counts are commonly considered standard for high-quality samples and proper imaging. However, less than ideal tissue, such as post-mortem human tissue, can contain age-related accumulations of highly autofluorescent molecules such as lipofuscin, AGEs, NADH, Flavins, and others [42]. The autofluorescence from these molecules may overwhelm and complicate single-molecule in situ signals, rendering data collection and analysis impossible in these situations. Additionally, post-mortem RNA degradation rapidly reduces the available in situ RNA targets, thereby reducing effective data throughput and underscoring the need for biochemical and computational methods that focus on comprehensive data collection [43].

The goal of this manuscript is three-fold. We explore the loss of information associated with axial sampling that exceeds the Nyquist-Shannon limit and provide a streamlined, GPU-accelerated computational approach to decoding iterative, barcoded RNA-FISH data. We additionally demonstrate a sample preparation method to reduce the deleterious effects of autofluorescence in non-ideal samples.

## RESULTS

### MERFISH decoding approach

A key component of any imaging-based spatial transcriptomic method is the computational imaging package used to turn the raw imaging data into localized RNA within segmented cells. One can frame the problem as an inverse problem, where the goal is to determine the most likely distribution of RNA 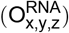 in continuous Cartesian space (x, y, z) given the observed images 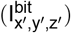 in a discrete space (x^*′*^, y^*′*^, z^*′*^) for each unique “bit”. The observed images are generated by an imaging system with numerical aperture (NA), excitation wavelength (λ_ex_), emission wavelength (λ_em_), point spread function (PSF), camera pixel size (Δx, Δy), discrete axial spacing (z^*′*^), camera gain (g_cam_), camera offset (offset_cam_), and excitation illumination profile (illum(λ_ex_, x, y, z)). We additionally require knowledge of the encoding strategy 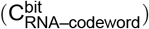 that connects individual iterative observations (bits) to the encoding-readout probe hybridization scheme (RNA-codeword) as well as image registration using fiducial markers, and any other prior information available, such as the Hamming weight and distance for Viterbi decoding [10, 41] or parity matrix for Syndrome decoding [13]. For Viterbi decoding, we seek to invert the forward imaging model,

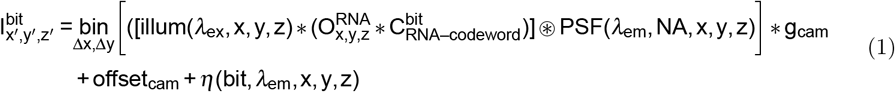

to solve for 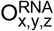. However, inverting Equation (1) to solve for 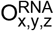 is confounded by the blurring operation introduced by the system PSF, the finite sampling of the camera Δx, Δy that locally bins pixels from continuous space (x, y) to a discrete space (x^*′*^, y^*′*^), the discrete axial sampling z^*′*^, and finally η, an unknown noise factor that includes instrument noise and additional sources of “noise” originating from biochemical reactions. These reactions include binding poly-adenylated (polyA) RNA to the gel matrix, encoding probes binding to RNA, readout probes binding to encoding probes, efficiency of “erasing” the readout probes in between readout rounds, photobleaching, and other unknown reactions.

We designed `merfish3d-analysis` to efficiently analyze non-axial Nyquist-Shannon sampled (called 2D from here on out) and approximately axial Nyquist-Shannon sampled (called 3D from here on out) iterative RNA-FISH experiments, including iterative smFISH and MER-FISH. The software performs the necessary data conversion and processing steps (camera correction, parameter-free deconvolution, linear and non-linear data registration, neural network spot prediction, self-supervision decoding parameter optimization, decoding, and filtering) on a CUDA-enabled workstation using a mixture of Python scientific processing libraries and CUDA kernels (Methods). Building on previous work, we include as many “self-supervised” steps as possible to limit the number of required user parameters. For deconvolution, we implemented a tiled, Andrew-Biggs accelerated version of gradient consensus deconvolution on the GPU. Using this approach, the data quality automatically determines the optimal number of deconvolution iterations, and images larger than the GPU memory can be deconvolved. To ensure that the pixel decoding only applies to pixels that contain RNA spots, we modified an existing neural network spot predictor (U-FISH [44]) to provide probability maps that are >0 where there is a potential spot-like feature in the deconvolved data and =0 otherwise. Finally, to optimize the bit normalization values that convert pixels potentially containing MERFISH bits to the range [0,1] for “soft” Viterbi decoding, we implemented an iterative approach. This approach begins with a guess based on all pixels and iteratively converges to bit normalizations by decoding a subset of tiles. The package automatically scales for multiple GPUs and leverages a modern, highly compressed, chunked file storage strategy for efficient data input and output. Here, we demonstrate the performance of the package on a series of simulated and real datasets.

### MERFISH simulations

To begin, we verified the accuracy of `merfish3d-analysis` and evaluated the impact of sub-sampling in the axial dimension by analyzing a series of realistic MERFISH simulations. We used a 16-bit Hamming Weight 4 Distance 4 codebook to encode codewords for 120 RNA species, along with 20 blank controls. We simulated two different spatial distributions of ground truth RNA molecule locations to test the effect of crowding. In the “uniform” condition, we randomly distributed RNA molecules throughout the simulation volume. In the “cell” condition, we primarily randomly distributed RNA molecules inside 3D cylinders with the typical volume of a cell (*∼*10 µm diameter). Table 1 summarizes the simulated RNA molecule numbers and densities.

**Table 1.** MERFISH simulation parameters. Simulation parameters for MERFISH simulations

|  | Uniform | Cells |
| --- | --- | --- |
| Number of RNA species | 120 | 120 |
| Total RNA molecules | 1200 | 400 in cells; 200 outside |
| Number of molecules per RNA species | $\sim 9.2$ | $\sim 9.2$ |
| Fluorescent spot density (molecule per $\mu\text{m}^2$) | $\sim 5 \cdot 10^{-2}$ | $\sim 4 \cdot 10^{-2}$ in cells |

For each of the ground truth simulations, we generated synthetic microscopy data at three different axial spacings: 0.315 µm, 1.0 µm and 1.5 µm. For the synthetic microscope, we used an objective with magnification of 100 *×* and numerical aperture of 1.4; tube lens of focal length 200 mm; a camera with pixel size of 6.5 µm, gain of 2 ADUs/electron, and offset of 100 ADUs; fiducial marker emission wavelength of 0.520 µm; and MERFISH readout dye emission wavelengths of 0.590 µm and 0.670 µm.

We simulated diffraction-limited fiducial markers and two bits of the codebook over eight imaging rounds using a detailed, probabilistic framework using realistic biochemical and measurement noise models (Methods, Supplemental Section 1).

### MERFISH simulation analysis

For all simulations, we ran the standard `merfish3d-analysis` pipeline, performing 3D decoding for axial spacings of 0.315 µm and 2D decoding for axial spacings of 1.0 µm and 1.5 µm. The results are summarized in Table 2 and Figures 1, 2. For both spatial simulation types, we found near perfect recovery of the location and identity of the ground truth RNA for 0.315 µm axial spacing. We saw reasonable recovery for 1.0 µm axial spacing and a loss of *∼* 35 % of RNA molecules for 1.5 µm axial spacing.

**Table 2.** Summary statistics of decoding results for MERFISH simulations. Decoding results for MERFISH simulation of randomly distributed RNA molecules (Uniform MERFISH) and MERFISH simulation of RNA molecules within cells (Cells MERFISH) at three different z spacings 0.315 µm, 1.0 µm and 1.5 µm using merfish3d-analysis with the default parameters. Precision, Recall, and F1 scores are calculated using a greedy matching algorithm with a search radius of 1.0 µm. All values rounded to two significant digits.

| Simulation Type | Axial Spacing | Precision | Recall | F1 score | Figure |
| --- | --- | --- | --- | --- | --- |
| Uniform MERFISH | $0.315\mu\text{m}$ | 1.0 | 1.0 | 1.0 | 1 |
| Uniform MERFISH | $1.0\mu\text{m}$ | 0.93 | 0.92 | 0.93 | 1 |
| Uniform MERFISH | $1.5\mu\text{m}$ | 0.66 | 0.65 | 0.66 | 1 |
| Cells MERFISH | $0.315\mu\text{m}$ | 1.0 | 1.0 | 1.0 | 2 |
| Cells MERFISH | $1.0\mu\text{m}$ | 0.96 | 1.0 | 0.98 | 2 |
| Cells MERFISH | $1.5\mu\text{m}$ | 0.7 | 0.62 | 0.66 | 2 |

**Figure 1.**
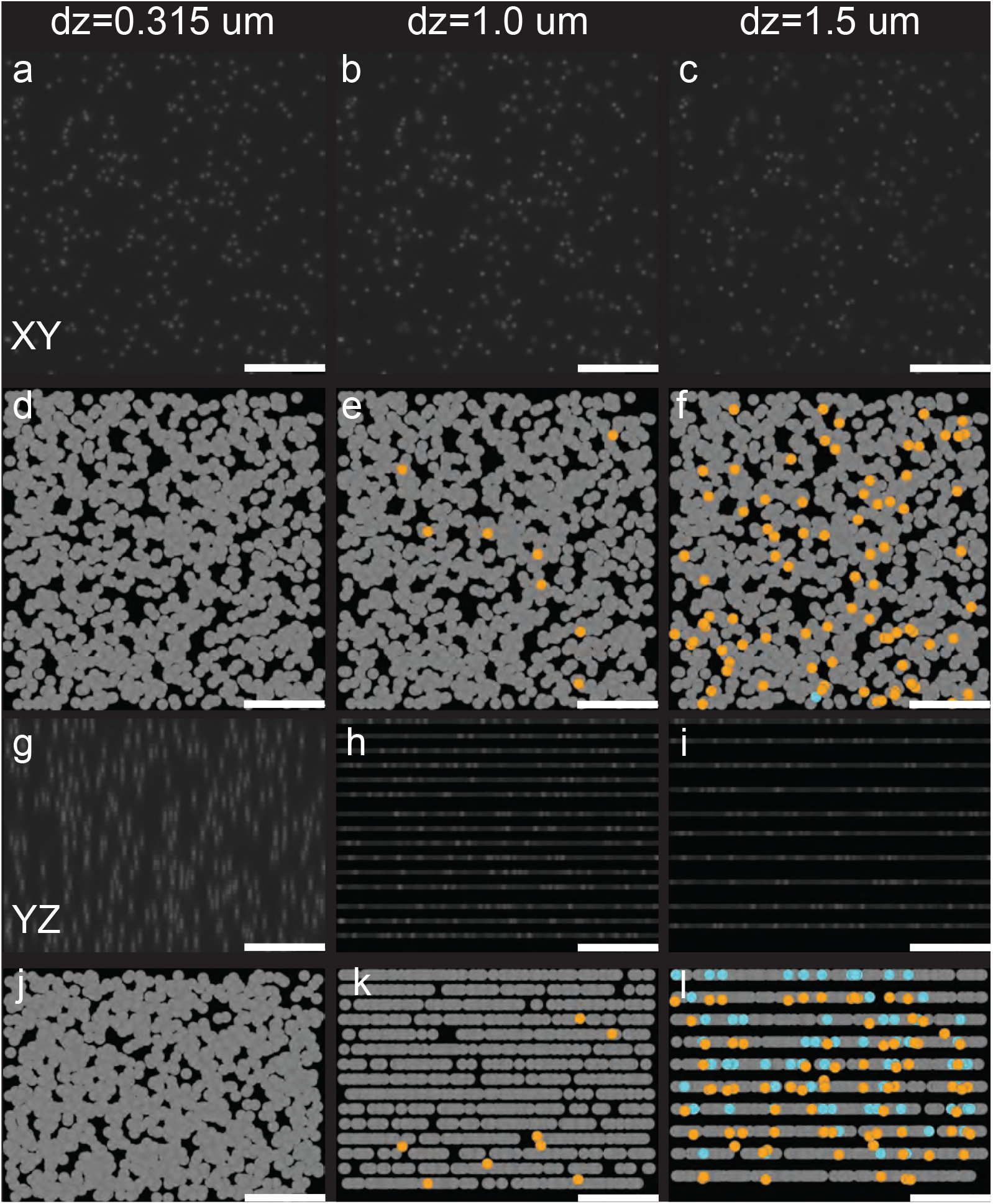
Uniformly distributed MERFISH simulation results. Axial sampling conditions for dz=0.315 µm, dz=1.0 µm, and dz=1.5 µm are organized by column. (a-c) Maximum Z projections (XY view) of one bit. (d-f) Maximum Z projections (XY view) of recovered RNA true positives (gray), false positives (cyan), and false negatives (orange). (g-i) Maximum X projections (YZ view) of one bit. (j-l) Maximum X projections (YZ view) of recovered RNA true positives (gray), false positives (cyan), and false negatives (orange). All scale bars 5 µm.

**Figure 2.**
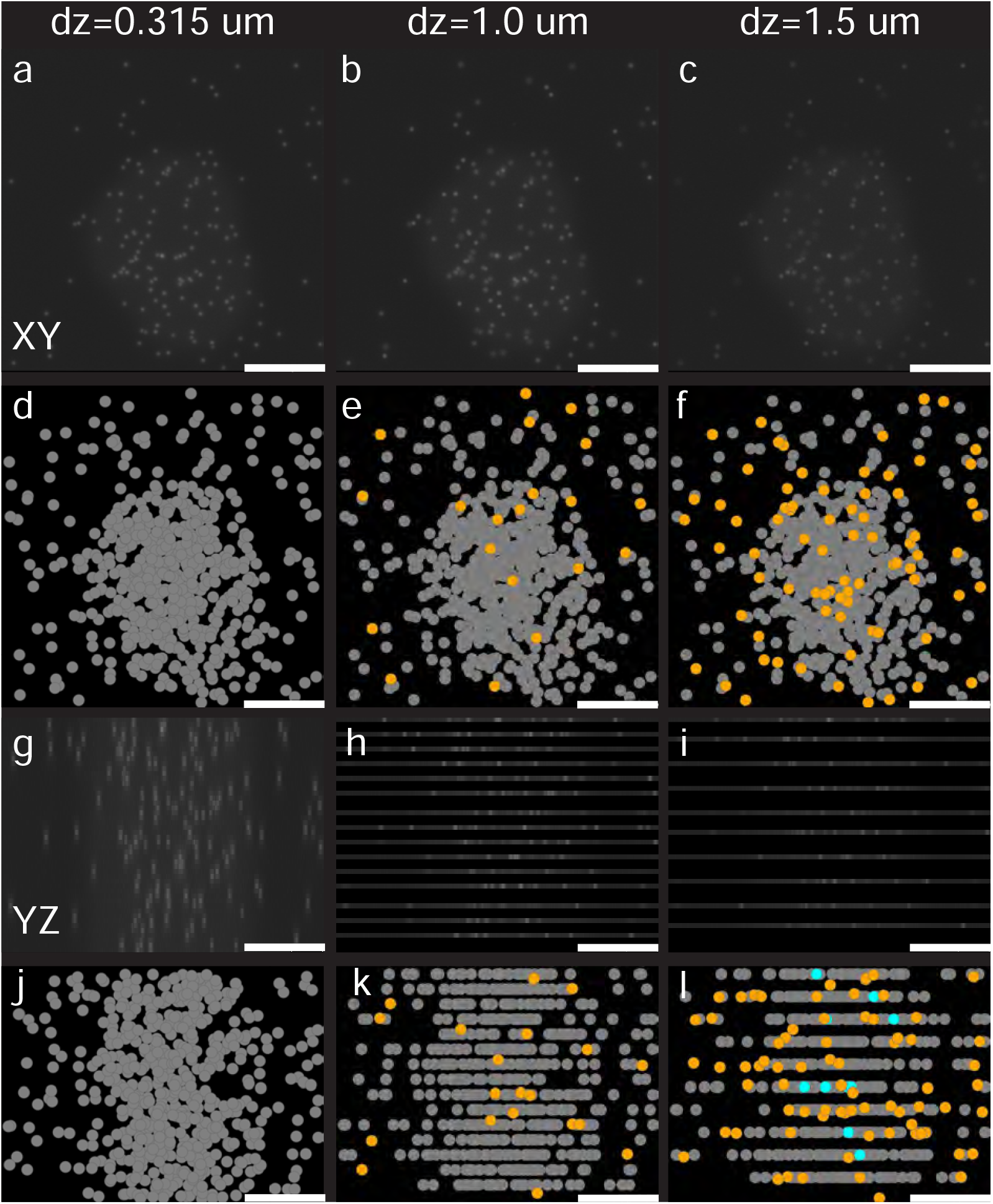
Cell distributed MERFISH simulation results. Axial sampling conditions for dz=0.315 µm, dz=1.0 µm, and dz=1.5 µm are organized by column. (a-c) Maximum Z projections (XY view) of one bit. (d-f) Maximum Z projections (XY view) of recovered RNA true positives (gray), false positives (cyan), and false negatives (orange). (g-i) Maximum X projections (YZ view) of one bit. (j-l) Maximum X projections (YZ view) of recovered RNA true positives (gray), false positives (cyan), and false negatives (orange). All scale bars 5 µm.

### Re-analysis of publicly available MERFISH data

After establishing good performance on simulated data, we proceeded to re-analyze a publicly available MERFISH dataset generated in the mouse motor cortex [45]. The motor cortex dataset was collected using a widefield microscope, with an axial spacing of 1.5 µm, over a depth of 9 µm. There are 447 FOVs within the dataset (Figure 3a). There are a few key differences in the final results between `merlin`, the package originally used to decode this data, and `merfish3d-analysis`. `merfish3d-analysis` deconvolution is self-guided and uses a vectorial model PSF, while `merlin` deconvolution uses a user-defined number of iterations and a 2D Gaussian approximation with a user-defined width. `merfish3d-analysis` predicts pixels that contain “spot-like” fluorescent features using a neural network, while `merlin` does not. Both packages perform pixel decoding, with `merlin` using deconvolved and then low-pass filtered data. At the same time, `merfish3d-analysis` uses deconvolved data weighted by the probability map produced from the neural network prediction.

**Figure 3.**
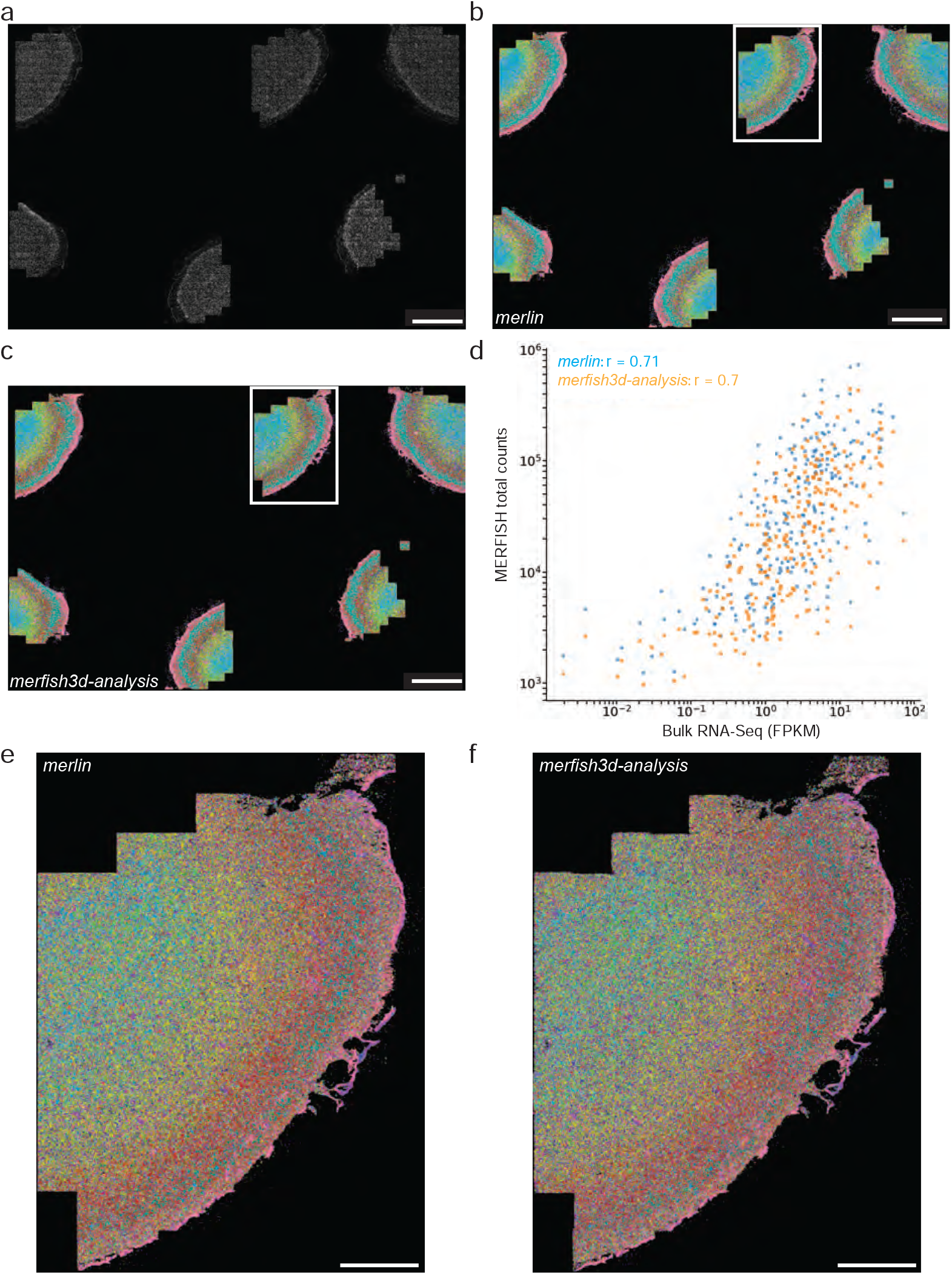
Publicly available mouse primary motor cortex MERFISH analysis. a) poly-dT overview of tissue slices in publicly available mouse motor cortex MERFISH dataset. b) `merlin` and (c) `merfish3d-analysis` decoding results, colored using Gini distance calculated for merlin data. d) Correlation between total MERFISH counts and bulk RNA-Seq FPKM. e) `merlin` and (f) `merfish3d-analysis` decoding results for highlighted region in (b) and (c), colored using Gini distance calculated for merlin data. Scale bars a-c) 1 mm e-f) 500 µm.

Figure 3b,c plots the provided RNA localizations, colored by Ginni distance clustering, for all six tissue slices in Figure 3a. For these tissue slices, the publicly uploaded `merlin` analysis contains *∼*17 million localized RNA. Because the original publication did provide the exact parameters and filtering values, we performed `merfish3d-analysis` decoding using default parameters and recovered *∼*11 million RNA. Despite this difference, we find a near identical correlation with bulk RNA-seq (using the same reference RNA-seq dataset from ref [45]) for both `merlin` and `merfish3d-analysis` decoding. We performed this analysis using all transcripts found within the highlighted tissue slice and only transcripts found with cells. Table 3 summarizes the correlation analysis. Similarly, the spatial Ginni clustering results are nearly identical between the two analysis approaches, with minimal differences across all tissue slices (Figure 3b,c,e,f).

**Table 3.** MERFISH vs bulk RNA-Seq Correlations. Pearson correlation metric for decoding results of mouse motor cortex MERFISH data. All values rounded to two significant digits.

| Package | Data type | Pearson coefficient |
| --- | --- | --- |
| `merlin` | All RNA within slice | 0.71 |
| `merlin` | Only RNA within cells | 0.7 |
| `merfish3d` | All RNA within slice | 0.7 |
| `merfish3d` | Only RNA within cells | 0.69 |

To understand the cause of the difference in total of number of RNA recovered, we manually inspected a few tiles to compare the decoding results of `merlin` and `merfish3d-analysis`. We discovered that most discrepancies are pixels where `merfish3d-analysis` calculations yield either no magnitude (i.e., there are no photons present across all bits) or the distance metric for those photons present is greater than a 1-bit error. In either case, `merfish3d-analysis` does not call an RNA at those pixels, but the uploaded `merlin` results contain an RNA molecule at that location (Supplemental Section 2).

### Analysis of post-mortem human olfactory bulb iterative RNA-FISH experiments

For less-than-ideal tissue types, specifically those containing highly autofluorescent endogenous and sometimes exogenous molecular features, we developed a sample preparation method. This method photochemically alters the sample using full-spectrum white light coupled with a UV filter to protect nucleic acid target molecules, in conjunction with clearing and proteolytic digestion. Figure 4a demonstrates the background autofluorescence for unperturbed, gel-embedded, 30 µm thick Human olfactory bulb tissue, with a baseline background autofluorescence ADU values of *∼*200 with peak feature grey values exceeding 1000, values that can overwhelm RNA FISH signals. A 60 minute clearing and proteolytic digestion reduces the autofluorescence signal (Figure 4b). Overnight sample photobleaching leads to a further reduction of background signal, approaching the camera offset of *∼*100 ADUs. (Figure 4c). Through the combination of gel embedding followed by chemical and physical bleaching, we were able to achieve high-quality, low autofluorescence staining of poly-dT anchor probes for entire 20 µm thick post-mortem human olfactory bulb samples (Figure 4d).

**Figure 4.**
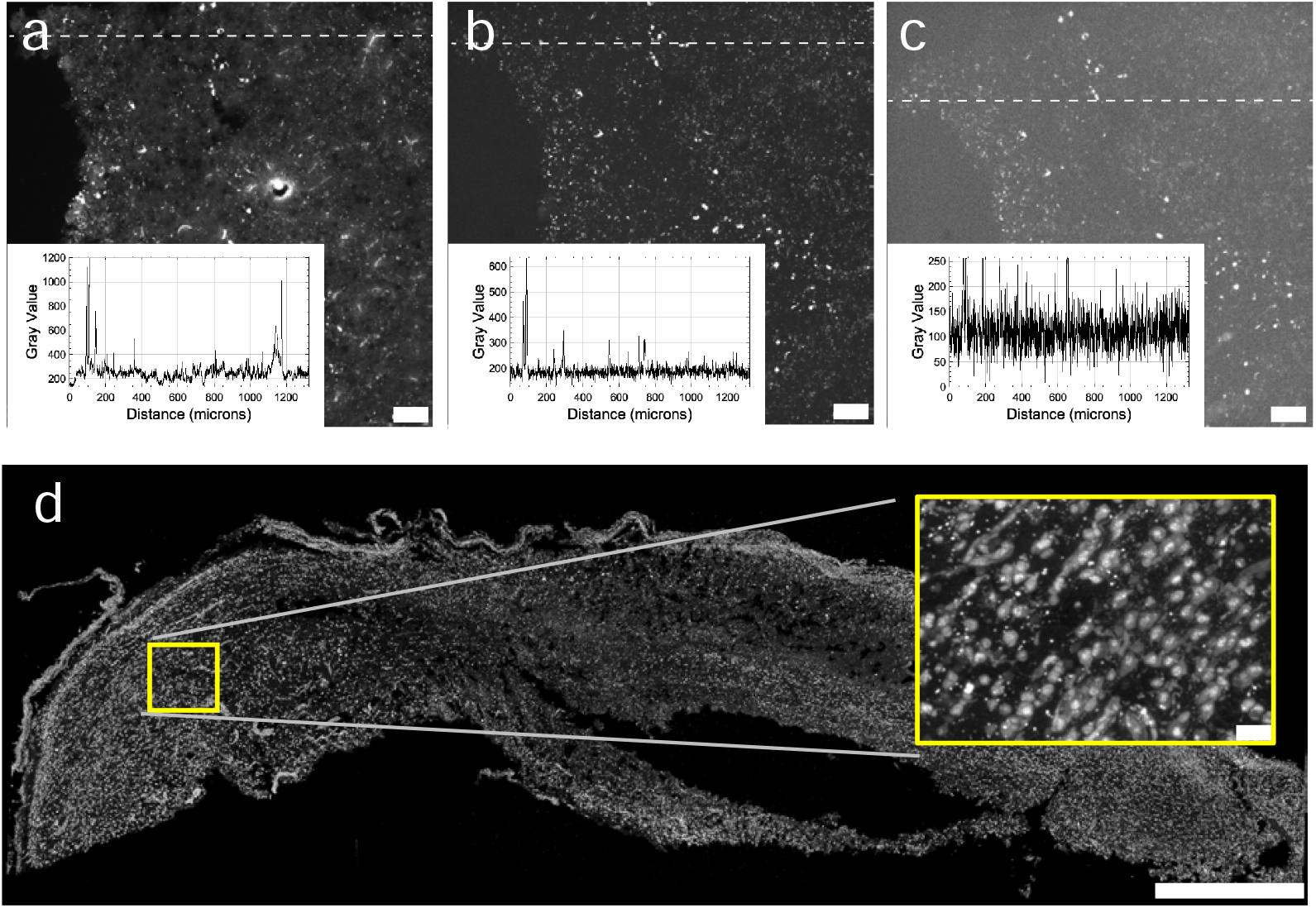
Chemical and physical photobleaching of aged tissue. a) Gel-embedded human olfactory bulb tissue imaged at 10*×* magnification, 470 nm excitation, and 50 ms exposure time. Inset: pixel intensity profile along the dotted line. b) Same sample as (a) re-imaged following a 90 min clearing and digestion using 4 % SDS clearing solution + 1 % proteinase k. Inset: pixel intensity profile along the dotted line. c) Same sample as (a) re-imaged following a 90 min clearing, digestion using 4 % SDS clearing solution + 1 % proteinase k, and overnight photobleaching. Inset: pixel intensity profile along the dotted line. d) Overview image of poly-dT probe for full OB slice after clearing, chemical treatment, and photobleaching. (a)-(c) scale bars, 100 µm); d) 1 mm

After performing the above photobleaching, we performed 3D MERFISH imaging on human olfactory bulb tissue using a 16-bit MERFISH 119-gene panel supplemented with two sequential smFISH readouts (codebook and panel design in Supplementary File 2) using a homebuilt widefield microscope and microfluidic controller. The olfactory system in mice is well studied, and in situ RNA measurements, including MERFISH, have found that each olfactory receptor gene is commonly clustered around two glomeruli structures (refs). However, little to no information is available on the molecular and cellular distribution of olfactory receptors in the human olfactory bulb.

The tissue morphology (poly-dT, two smFISH readouts) and spatial extent of the experiment ROI are shown in Figure 5a,b. During imaging, we used an axial sampling of 0.315 µm over a depth of *∼*15 µm and 70 slightly overlapping FOVs within the ROI. The results of MERFISH decoding (Figure 5c) found evidence of clustering for two olfactory receptor RNAs, OR2A20P and OR3A4P, both of which are co-localized with a synaptic marker (SVB2) (Figure 5d).

**Figure 5.**
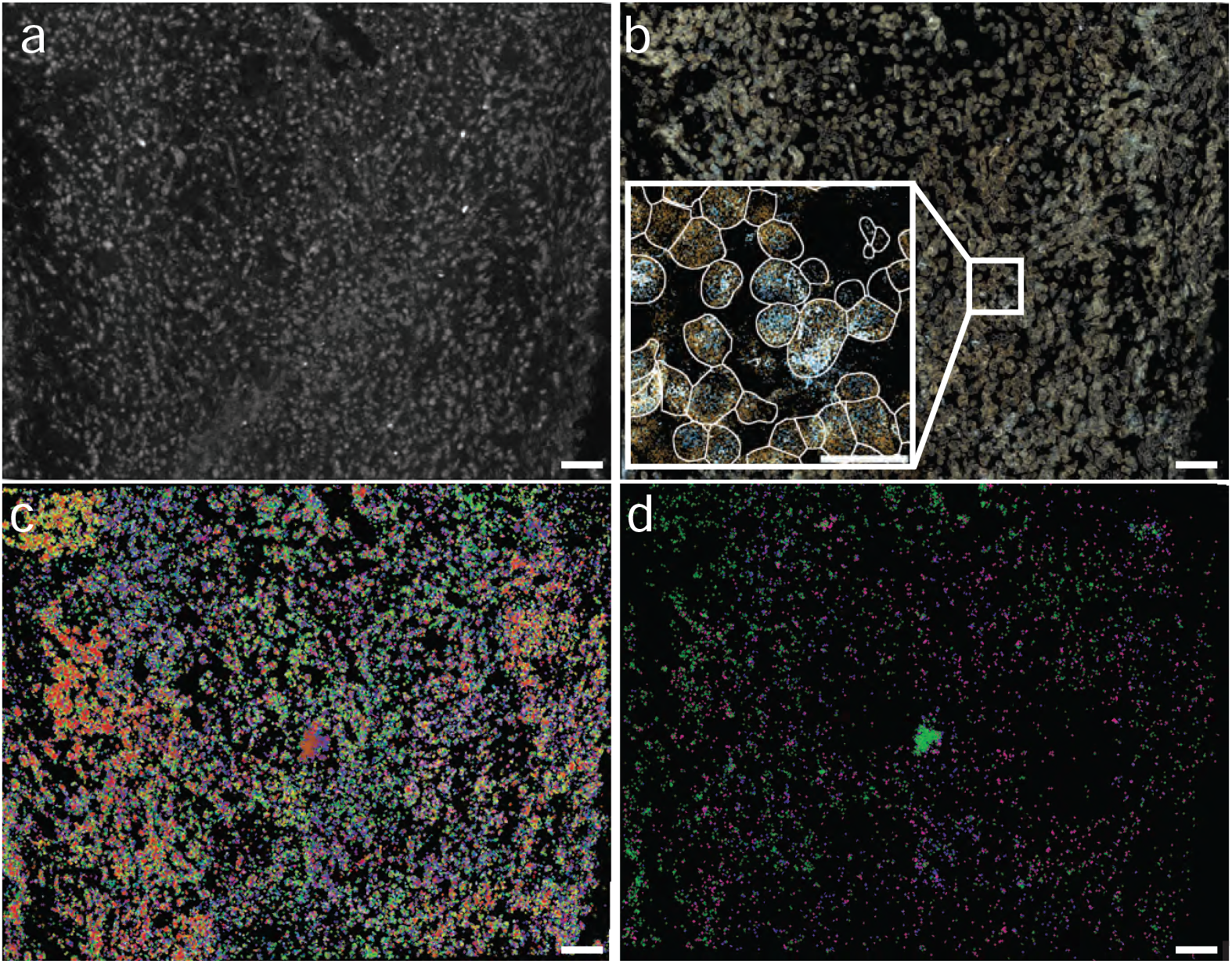
Human olfactory bulb MERFISH experiment and analysis. a) Maximum Z projection of poly-dT signal (gray), b) Maximum Z projection of SYN2 (cyan) and NCAM1 (orange) smFISH readouts overlaid on cell outlines (white). c) Maximum Z projection of Gini distance colored 16-bit MERFISH 119-gene panel data. d) Maximum Z projection for OR2A20P (green), OR3A4P (magneta) and SV2B (blue) RNA expression. All scale bars are 250 µm.

## DISCUSSION

The analysis of our simulations shows that for sparsely distributed RNA molecules, the impact of axial subsampling depends on RNA density. For uniform randomly distributed RNA at dz=0.315 µm and 1.0 µm, we found an F1 score of 1.0. As dz increases, the recall score decreases, indicating a loss in recovered RNA solely due to the change in axial sampling strategy. By increasing the density of the simulation, we found that the recovery of RNA molecules rapidly drops off as a function of increasing dz. However, precision in the recovered spots remains high. As the likelihood of two fluorescent-labeled RNAs being in close spatial proximity increases, the uncertainty in the decoding procedure exceeds acceptable levels, leading to decoding failures. By relaxing the magnitude threshold and minimum “spot” size thresholds, it is possible to finetune the pixel decoding results to obtain improved results. However, for real experiments where the ground truth is not known, blind tuning of the decoding parameters will eventually lead to a higher number of false positives (Supplemental Section 2). Ultimately, we suggest that the desired experimental outcomes should guide the choice of lateral and axial sampling, as well as the number of tiles acquired. If the goal is to survey the RNA molecules within a sample accurately, then spatial sampling near the axial Nyquist-Shannon spacing is essential. If the goal is to survey molecules present in a larger volume of tissue, relaxing axial sampling can still provide a higher capture rate than traditional spatial sequencing-based methods.

Moving to real data, there are several data formatting and experimental realities we sought to address, which were not present in our simulations. Many of the existing iterative RNA-FISH analysis packages are hard-coded to the experimental format [13, 37, 41, 46], sacrifice performance or have a large number of required user input parameters [29], or do not provide data conversion at all [10, 30, 33–35]. By defining a structured high-performance datastore format, we aimed to enable the conversion and analysis of iterative RNA-FISH experiments acquired using any instrumentation. To support this, we demonstrate the conversion and analysis of simulations, existing public data, and data from our own instruments, each with a unique file format.

By implementing all algorithms using lazy GPU computing with chunked processing, we aimed to enable high-performance analysis on local workstations. Many of the packages intended for analyzing larger, tiled datasets rely on computing cluster environments to achieve sufficient parallelism for reasonable run times. As large iterative RNA-FISH datasets can exceed tens to hundreds of terabytes, transferring this data to a remote cluster is a costly exercise and a barrier for many researchers. On a standard workstation with two GPUs, we converted and processed the 448-tile, 38-bit public (22-bit MERFISH + 16 smFISH bits) dataset in Figure 3 in roughly 60 hours.

The modular nature of `merfish3d-analysis` allows for easy upgrades while not changing the public API for the end user. For example, during development, we changed our deconvolution approach from the `pycudadecon` package [47] to our own custom GPU implementation of gradient consensus deconvolution [48]. This change did not require reworking the entire processing package; it only involved replacing the deconvolution module while keeping the input and output variables consistent. The advantage is that gradient consensus deconvolution is selfsupervised, eliminating the need for the user to set the number of iterations or a regularization strength. We anticipate that as new neural network advancements become available, we will continue to upgrade `merfish3d-analysis` to take advantage of the increased computational speed or accuracy.

Autofluorescence and target molecule degradation in non-ideal tissues complicate FISH signal strength and increase off-target labeling. We demonstrated that sample photo-bleaching reduces auto-fluorescence in both background and peak intensity features, thus improving the acquisition signal-to-background ratio. Reducing the background is particularly important for short-sequence or high-homology RNA in situ targets, where relatively few high-stringency probes are available. The designed sample pre-treatment resulted in an improved poly-dT signal for the post-mortem human olfactory bulb samples we tested, suggesting that much of the poly-adenylated RNA remained in the gel with minimal autofluorescence. However, after performing multiple MERFISH experiments, we found significantly lower RNA counts per cell in the post-mortem human tissue used in this study when compared to tissue sourced from animals. New methods for RNA preservation and potentially recovery of partially degraded transcripts would help open imaging spatial transcriptomics to a broader variety of archival samples. Despite the lower RNA per cell counts, we did find evidence of high expression of 2 olfactory receptor pseudogenes (OR2A20P and OR3A4P) out of the 55 we probed for. Complicating this analysis is the high homology between OR2A20P and OR3A4P, with a nearly 50 % global sequence alignment based on NBCI reference sequences. Further studies are required to determine if the high homology across human olfactory receptor genes and pseudogenes will limit our ability to probe for specific receptors. One strategy may be to use a “pan” olfactory receptor FISH probe set for high homology sequences, combined with high stringency probe selection for unique sequences.

Moving forward, we aim to reduce further the time associated with file reading and writing by enhancing parallelism during the conversion from existing experiments and by directly writing the `merfish3d-analysis` datastore format from our in-house microscopes. Another area for improvement is to create a streaming interface that allows users to transfer or process data on the fly during an experiment, enabling as much processing as possible before the entire dataset is collected. Unfortunately, because each imaging round must be registered to a standard reference coordinate system, decoding cannot occur until all data is acquired.

Another target for optimization is the self-supervised bit normalization algorithm. While iteratively decoding and using only RNA-associated codewords to extract the bit offset and bit normalization values is a robust approach, we can likely remove this process entirely by training a neural network to transform preprocessed MERFISH data directly to the correct values. Such an improvement would help further streamline the decoding and reduce the overall processing time.

We additionally tested `merfish3d-analysis` on simulated iterative smFISH data (Supplemental Section 2), with no error correcting rounds. We achieved reasonable decoding results after empirically tuning the pixel decoder parameters, suggesting that it may be possible to extend pixel-decoding to iterative smFISH experiments. While such experiments lack the multiplexing capability of barcoded codebooks, it is possible to target high-expression genes or genes with a low total FISH probe number using iterative smFISH.

In summary, we have introduced a new software package, `merfish3d-analysis`, that performs all the steps necessary to convert raw microscope data into decoded RNA locations for iterative RNA-FISH experiments. In this manuscript, we have focused on one specific iterative method, multiplexed error-resistant RNA-FISH (MERFISH) [10]. However, users can apply the work here to any iterative method that requires decoding fluorescence intensity values acquired over multiple chemistry rounds using a known codebook. `merfish3d-analysis` is fully GPU-accelerated and aims to minimize the number of user-tunable parameters as much as possible through self-supervised algorithms. We demonstrated the performance of `merfish-analysis` on simulated data with known RNA locations, re-analysis of publicly available data, and analysis of post-mortem human olfactory bulb tissue.

## METHODS

### MERFISH simulations

We developed a probabilistic framework to simulate the forward model for MERFISH images for randomly distributed RNA with realistic biochemical, photon, and measurement noise. Details are provided in Supplemental Section 1.

### GPU-accelerated image processing and decoding

All processing used `merfish3d-analysis` v0.7.6, a lazy, chunked, end-to-end MERFISH analysis package with multi-Nvidia GPU support.

#### Data format

Raw microscope data were converted to a `zarr v2` file format organizing fiducial versus readout channels across tiles and rounds. The `tensorstore` Python package handled disk I/O. Known transformations between stage (x,y,z) axes and camera axes were applied during processing to ensure downstream analysis proceeds in a corrected global coordinate system.

#### Local-coordinates image processing

Camera ADU to photon conversion. Camera ADU counts were converted to photons using the manufacturer supplied conversion factor, offset, and quantum efficiency curves.

##### Illumination correct

Flatfield correction images were calculated from camera corrected data for each “channel”. For all independent channels, 100 tiles were randomly selected both within space and across all iterative imaging rounds. The illumination image was estimated from the randomly selected tiles using the python implementation of BaSiC [49].

##### Self-supervised deconvolution

All camera and illumination corrected images were deconvolved using a GPU-enabled gradient consensus Richardson–Lucy algorithm with Andrew-Biggs acceleration [48]. For non-Nyquist-Shannon sampled 2D data, a theoretical in-focus 2D point spread function (PSF) was used; for Nyquist-Shannon sampled 3D data, a theoretical 3D PSF was used. PSFs were computed from microscope parameters using a vectorial model [50]. Volumes exceeding GPU memory were processed in chunks using custom strategies and `ryomen` [51].

##### Registration using fiducial channel

Within each tile, deconvolved fiducial images were registered across fluidics rounds using two steps: (i) full 3D cross-correlation on downsampled data to estimate coarse (x,y,z) translation; and (ii) discrete deformable image registration to account for non-rigid translation [52]. The rigid and non-rigid transforms were used to map both fiducial and readout channels for each tile/round back to the initial fiducial round.

##### Self-supervised pixel content prediction

Deconvolved, registered readout images were passed through a modified U-FISH spot-prediction framework to estimate the probability that a pixel contains fluorescence from an RNA-FISH spot [44]. Pixel-wise probability maps were saved for use in decoding.

##### Pixel-based decoding

Within a given tile, deconvolved data and U-FISH predictions were loaded for all readout bits. Raw data were scaled from [0, 1] per bit:

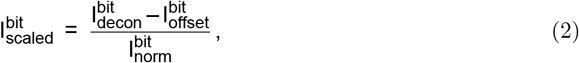

and low-pass filtered (2D kernel in each z plane for non-Nyquist-Shannon sampled; 3D kernel for Nyquist-Shannon sampled):

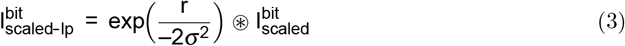

Pixels were gated by U-FISH predictions with a threshold (commonly 0.2):

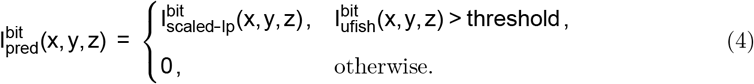

For each voxel, we computed the L2 norm across bits:

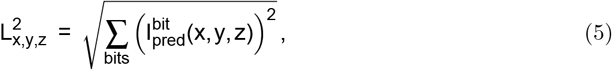

and determined the codeword minimizing the Euclidean distance between the normalized pixel vector and the codebook:

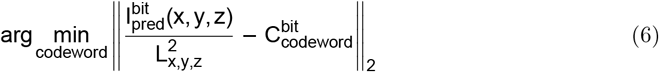

##### MERFISH pixel decoding and spot calling

Spatially connected pixels were aggregated into spots for RNA decoding: 2D connected components for non-Nyquist-Shannon sampled data and 3D for Nyquist-Shannon sampled data. Decoded RNA was accepted only if it had normalized intensity magnitude within the one-bit error window, ≥1.1 and ≤2.0, for properly normalized Hamming weight-4, distance-4 codebook) and Euclidean distance less than the one-bit error radius (0.5176 for Hamming weight-4, distance-4). Labeled RNA species were then warped to global coordinates using previously calculated affine transforms.

##### Iterative smFISH pixel decoding and spot calling

Spatially connected pixels were aggregated into spots for labeling as one of the 10 known good codewords, using 2D connected components for non-Nyquist-Shannon sampled data and 3D for Nyquist-Shannon sampled data. Spots only passed thresholding for labeling if they had a normalized intensity magnitude within the range ≥0.75 and ≤1.75 and a Euclidean distance less than 1.0. Labeled RNA species were then warped to global coordinates using previously calculated affine transforms.

##### Self-supervised bit normalization optimization

To scale data on [0, 1], initial global normalization (foreground) and offset (background) parameters were estimated using the 95th and 5th percentiles across N random tiles per bit. From these, new sets of M random tiles were decoded iteratively to update the bit foreground/background parameters. For all accepted non-blank codewords, the median foreground normalization was computed over on-bits and the median background over off-bits per iteration. Convergence typically occurred in fewer than 10 rounds.

##### False-discovery filtering

To limit false discovery, a logistic regression classifier was trained on gene-encoding and blank-encoding codewords. Codewords were filtered until the false discovery rate (FDR) fell below a user-defined threshold (commonly 0.05):

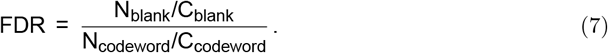

Class balancing utilities in scikit-learn were used to subsample blanks for adequate training (cite).

##### Global coordinate system assignment

Using known tile affine transforms and voxel size, spots were transformed from local pixel to world coordinates. Spots were then filtered to remove duplicates: 3D duplicates in tile overlaps for 3D data, and both tile-overlap and adjacent-plane duplicates for 2D data.

#### Global-coordinates image processing

Registration using first fiducial round. Deconvolved fiducial images from the first fluidics round were registered across tiles using `multiview-stitcher` [53]. The known stage-to-camera affine transform was applied to each tile; 3D cross-correlations were computed for overlapping tiles; and the global configuration minimizing tile dispersion was solved on downsampled 3D data. Fused downsampled 3D volumes and Z-maximum-projection 2D overviews were generated for initial cell segmentation.

##### Cell segmentation using poly-dT data

The fused Z-maximum projection 2D overview was segmented using `Cellpose-SAM` [54]. Parameters were optimized on a subset of tiles via GUI, then applied programmatically. The resulting 2D masks were converted to pixel-coordinate outlines, warped to global coordinates, and saved as ImageJ ROI files for downstream use.

##### Cell assignment using segmentation masks

Each 2D outline was extruded through Z to form a cylinder spanning all planes within the mask. RNA within a mask were assigned that cell’s identifier. RNA outside any mask were assigned –1 to indicate extracellular localization.

##### Visualization

poly-dT overviews, poly-dT with cell boundaries, and individual tiles were visualized in `FIJI` [55]. Scatter plots of RNA molecules were generated using `fishSCALE` [41]. For MERFISH plots, genes were sorted by Gini clustering index and colored using a rotating colormap to highlight spatially similar expression.

### Photobleaching study

Olfactory bulb samples were procured through Anabios as described in “Sample Procurement” and sagittally sliced on a cryostat to 30 µm-thick slices. The experimental design of this photobleaching study follows the iterative smFISH protocol as described above including tissue slicing, mounting, gel embedding and clearing. Poly-adenylated mRNAs were anchored to the gel matrix using the poly-dT LNA anchor probe (linker) to retain RNA through the clearing process. Encoding probe staining was not included. Samples were gel-embedded and imaged prior to clearing to capture naive endogenous tissue autofluorescence. Imaging was performed on a Nikon Ti-Eclipse at 10 *×* magnification at 470 nm excitation at 50 ms exposure. Samples were then cleared for 60 minutes in 4 % SDS clearing solution supplemented with 1 % proteinase then rinsed with preheated 2 *×* SSC Wash buffer 3-4 *×* for 5 minutes each prior to re-imaging of the same area. Finally, samples immersed in 5 *×* SSC buffer with 3 mg mL^−1^ PVSA or 1 % Murine RNAse inhibitor, were photobleached overnight for > 12 hours in a custom cooled chamber with a white light setup and a UV filter used to protect the sample from ultraviolet radiation. Once photobleached, samples were rinsed in 2*×* SSC buffer 3-4*×* for 5-10 minutes each. The same region of the sample was then re-imaged.

### Human olfactory bulb MERFISH panel design

#### 119-gene human encoding probe library design

A list of 2380 unique MERFISH encoding probes were designed using the open-source software PaintSHOP [56] against the hg38 (isoform-flattened) genome targeting 119 genes. Encoding probe template sequences were purchased as a single-stranded oligo library (IDT, Inc.; pooled ssDNA oligos).

Encoding probes feature a target region complementary to the RNA species of interest, bridge sequences complementary to readout probes, and priming regions used to construct working encoding probe stocks from the purchased oligo library. Bridge sequences were derived from ref [10, 19] as the reverse complement of the described bDNA readout probe sequences.

For PCR amplification and reverse transcription, we used the following 20-nt primer sequences: forward CTAGGAGTCTAGCTACTACC and reverse TAATACGACTCACTATAGGG. Twenty probes per RNA target species were designed in PaintSHOP using the “unify number” function and default advanced settings (maximum off-target score = 200, maximum k-mer count = 5, minimum prob value = 0).

For 16-bit MHD4 barcoded experiments, four bits are assigned per RNA species whereby PaintSHOP assigns three of the four possible bridge sequences to each encoding probe. Priming regions used in probe synthesis (PCR, in-vitro transcription, and reverse transcription) were appended to the terminal 5’ and 3’ ends of the encoding probes. To construct the encoding probe oligo library in PaintSHOP, a list of candidate gene identifiers, bridge sequences, and PCR priming regions were compiled and appended using the application.

#### Two-gene smFISH encoding probe design

In addition to the 119-gene MERFISH panel, a set of smFISH probes were designed for two glomerular marker genes, SYN2 and NCAM1, to highlight glomeruli in the olfactory bulb. smFISH probes were designed in PaintSHOP using the RNA probe design function and the hg38 genome. Using accession numbers for these genes, we constructed lists of 66 and 104 probes for SYN2 and NCAM1, respectively. Two alternate bridge sequences were appended together with the same 20-nt priming sequences used in the MERFISH probe set design. smFISH probes were combined with the MERFISH probe set and ordered together as a single oligo pool from IDT, Inc.

#### Encoding probe panel construction

Encoding probes were constructed from the oligo library sequences following previously described in refs. [17, 18, 20]. Briefly, the library was amplified using limited-cycle PCR with priming sequences CTAGGAGTCTAGCTACTACC on the 5’ end of each probe and CCCTATAGTGAGTCGTATTA (T7 reverse complement) on the 3’ end.

Using the PCR product as template, we performed in-vitro transcription (IVT), which simultaneously shortened probes by omitting the T7 promoter region. We then used reverse transcription with a modified, single uracil-incorporated reverse transcription primer (`CTAGGAGTCTAGCTACrUACC`) to generate cDNA. Uracil-Specific Excision Reagent (USER enzyme) was used to cleave uracil from the 5’ priming region, shortening the final probe lengths [18, 20]. All remaining RNA was digested by alkaline hydrolysis, and final single-stranded DNA probes were purified via column purification.

Probes were concentrated using ethanol precipitation or vacuum centrifugation and resuspended to *∼*500 µm in TE buffer (Life Technologies; Cat. AM9849).

### Sample preparation

#### Coverslip functionalization

For sample mounting, we functionalized 40 mm-diameter #1.5 coverslips (Bioptechs; Cat. 40-1313-0319) based on previously described cleaning, silanization, and poly-D-lysine (PDL) coating methods with modifications [17]. Briefly, coverslips were immersed in a 1:1 mix of 37 % HCl and methanol at room temperature for 30 min, rinsed three times in nuclease-free water, autoclaved in nuclease-free water, and rinsed once with 70 % ethanol.

Cleaned coverslips were maintained in 70 % ethanol until use. To silanize, cleaned coverslips were removed from ethanol and dried at 60 °C. Coverslips were immersed in a silanization solution consisting of 0.1 % (vol/vol) triethylamine (Millipore; Cat. TX1200) and 0.2 % (vol/vol) allyltrichlorosilane (Sigma; Cat. 107778) dissolved in chloroform for 30 min at room temperature, washed once with 100 % chloroform, and washed once with 100 % ethanol. Coverslips were baked at 60 °C for 2 h to fix the silane layer.

To improve sample adherence, we added an additional coating of 2 % aminopropyltriethoxysilane (APTES; Sigma; Cat. 440140) as previously described in ref. [46]. Coverslips were immersed in 2 % APTES in acetone with sonication for 2 min, followed by two rinses with nuclease-free water and one rinse in ethanol, then dried at 60 °C. Coverslips were then PDL-coated by immersion in a 0.1 mg mL^−1^ PDL solution (Sigma; Cat. SLBS8705) for 1 h at room temperature and rinsed three times with nuclease-free water. After drying, silanized and PDL-coated coverslips were stored at room temperature in a desiccation chamber [18].

#### Sample procurement

Human olfactory bulb tissue was procured through Anabios. Post-mortem tissue was surgically collected and immediately rinsed with ice-cold nuclease-free 1 *×* PBS containing 0.1 % murine RNase inhibitor (NEB; Cat. M0314L) to remove blood. Samples were incubated at 4 °C in 4 % paraformaldehyde in 1 *×* PBS for 24 h, rinsed in 1 *×* PBS, then cryoprotected by immersion in a 15 % and then 30 % RNase-free sucrose series (in 1 *×* PBS) before embedding and freezing in OCT medium. Samples were stored at −80 °C.

#### Tissue slicing, processing, and permeabilization

Human olfactory bulb was sliced sagittally on a Leica model 1860 cryostat at −20 °C and transferred to functionalized coverslips. For this and all subsequent tissue processing steps, coverslip-mounted samples were maintained in RNase-pretreated 60 mm Pyrex petri dishes with lid.

Tissue slices were post-fixed to the coverslip surface with 4 % PFA for 15 min at room temperature and rinsed with 1 *×* PBS. To quench free aldehydes following fixation, samples were incubated for 5 min in 100 mm ammonium chloride and rinsed with 1 *×* PBS. To reduce heme and lipofuscin autofluorescence, samples were incubated in several drops of 3 % peroxide solution (ACD; Cat. 322380) for 10 min and rinsed twice with 1 *×* PBS. Slices were pretreated for 5 min in 4 % sodium dodecyl sulfate (SDS) clearing solution in 1 *×* PBS, then rinsed in 2 *×* SSC buffer four times for 5 min to remove residual clearing solution. Slices were permeabilized overnight in 70 % ethanol at 4 °C. Once permeabilized, slices were rehydrated in a 50 %, 30 % and 10 % ethanol series prior to encoding probe staining.

To minimize reagent use and confine buffers to the coverslip, a *∼*30 mm diameter hydrophobic barrier (Vector Laboratories ImmEDGE™ Hydrophobic Barrier Pen) was drawn around the sample and allowed to dry for 5 min. To prevent desiccation, the tissue was immersed in 1 *×* PBS during this time.

#### Probe staining procedure

Samples were first stained with an anchor probe prior to gel embedding/clearing/photobleaching, followed by encoding probe staining. Both anchor probe and encoding probe hybridizations used the same workflow.

Prior to staining, samples were pretreated with formamide wash buffer consisting of 40 % (vol/vol) formamide (Fisher; Cat. AM9342) with 1 % (vol/vol) Tween-20 (Sigma; Cat. P9416) in 2 *×* SSC buffer (Fisher; Cat. AM9763) for 30 min at 47 °C. Following pretreatment, samples were incubated in 125 µL hybridization buffer consisting of 40 % (vol/vol) formamide, 0.1 % (wt/vol) yeast tRNA (Life Technologies; Cat. 15401011), 1 % (vol/vol) murine RNase inhibitor (NEB; Cat. M0314L), 1 % (vol/vol) Tween-20, and 10 % (wt/vol) dextran sulfate (Sigma; Cat. D8906) in 2 *×* SSC plus probes. To minimize evaporation, an RNase-pretreated 25 mm diameter coverslip overlay was applied; the petri dish chamber was Parafilmed and placed in a humidified box inside a humidified hybridization oven.

Following hybridization, samples were incubated twice for 30 min at 47 °C in 40 % formamide wash buffer, then twice for 5 min at 47 °C in SSC-Tween consisting of 2 *×* SSC with 0.1 % Tween-20, and briefly rinsed twice with room-temperature 1 *×* PBS.

#### RNA anchor probe hybridization

To reduce autofluorescence, samples were gel embedded and cleared. Prior to gel embedding, to anchor and retain mRNA in the gel matrix, samples were stained with 1 µm poly-dT LNA anchor probe including a 20-nt readout sequence and a 5’ acrydite modification `/5Acryd/TTGAGTGGATGGAGTGTAAT T+TT+TT+TT+TT+TT+TT+TT+TT+TT+T`, as previously described [20], for 18 h to 24 h following the probe staining procedure above.

### Gel embedding, clearing, and photobleaching

Gel embedding and clearing followed established protocols [17]. Anchor probe–stained samples were incubated in degassed acrylamide monomer solution consisting of 4 % (vol/vol) 19:1 acrylamide:bis-acrylamide (BioRad; Cat. 1610144), 60 mm Tris-HCl (Fisher; Cat. AM9856), and 0.3 m NaCl (Fisher; Cat. AM9753) for 30 min. Samples were equilibrated for 2 min in acrylamide gel solution consisting of the monomer solution with polymerizing additives 0.03 % (wt/vol) ammonium persulfate (Sigma; Cat. A3678) and 0.15 % (vol/vol) TEMED (Sigma; Cat. T9281), and fiducial YG beads (PolySciences; Cat. 17150-10) at *∼* 10^−3^ relative to stock concentration. The gel solution was refreshed, covered with a gel-slick–pretreated 25 mm overlay coverslip, and polymerized for 1.5 h at room temperature. To maintain a thin gel, necessary for optimal probe diffusion, excess gel solution was pipetted or wicked away from the sides of the overlay coverslip.

For photobleaching, samples were immersed in room-temperature 5 *×* SSC storage buffer supplemented with 1 % murine RNase inhibitor or 3 mg mL^−1^ poly(vinylsulfonic acid, sodium salt) (Sigma; Cat. 278424) and overlaid with a UV filter (Thorlabs; Cat. FGL435S). The chamber was shielded with foil, leaving only the UV filter exposed. The photobleaching setup consisted of a (6 in *×* 6 in), 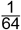 in-thick lidless metal-framed box, a tablet semiconductor cooler affixed underneath, and a 36 W full-spectrum grow light overhead. Samples were photobleached overnight (12 h to 18 h typically).

After photobleaching, samples were rinsed with 1 *×* PBS then incubated at 37 °C in preheated SDS clearing solution consisting of 4 % SDS (Sigma; Cat. 75746) in 1 *×* PBS supplemented with 1 % proteinase K (NEB; Cat. P8107S) for 18 h to 24 h. Cleared samples were rinsed by incubating in 37 °C 2 *×* SSC four times for 5 min to 15 min each.

#### Encoding probe hybridization

MERFISH encoding probe staining followed previous methods with minor modifications [18, 56]. A detailed protocol is provided online [57]. Gel-embedded, photobleached, and cleared samples were stained with encoding probes at 1 nm per encoding probe following the staining procedure above and incubated in hybridization buffer for 48 h to 72 h.

### Iterative readout staining using automated fluidics exchange

Iterative readout staining followed previously described methods with an alternate imaging buffer [18]. Coverslip-mounted samples were placed into a Bioptechs FCS-2 flow cell chamber (Bioptechs; Cat. 03060319-2-NH) with a 14 mm*×*24 mm, 0.75 mm-thick rectangular gasket (Bioptechs; Cat. 1907-1422-750) and connected to a custom, peristaltic pump–driven automated fluidics system. This system was adapted from a Moffitt Lab design [21].

Fluidics buffers included: readout hybridization buffer (RHB) consisting of 2 *×* SSC with 10 % (vol/vol) ethylene carbonate and either 0.1 % (vol/vol) murine RNase inhibitor or 3 mg mL^−1^ poly(vinylsulfonic acid, sodium salt) (Sigma; Cat. 278424) plus 3 nm of each appropriate readout probe; readout wash buffer (RWB) consisting of 2 *×* SSC with 10 % (vol/vol) ethylene carbonate and 3 mg mL^−1^ poly(vinylsulfonic acid, sodium salt); 2 *×* SSC; cleavage buffer (CB) consisting of 50 mm tris(2-carboxyethyl)phosphine (TCEP) in 2 *×* SSC; and imaging buffer (IB) consisting of degassed 50 µm Trolox-quinone, 50 µm 3,4-dihydroxybenzoic acid (PCA), 5 µm Trolox, and 50 µm NaOH in 2 *×* SSC supplemented with 2 µL mL^−1^ rPCO enzyme (OYC Americas; Cat. 46852004) immediately before use. All automated fluidics were at room temperature.

Prior to connecting the sample to the fluidics, buffers were primed to the valve positioner and the system was rinsed with 2 *×* SSC to remove air bubbles. A bubble trap (Darwin Microfluidics; Cat. VF-KBT-M-A) with applied negative pressure was added to capture any bubbles during the experiment.

Each readout round consisted of sequential multiplexed readout stainings (two paired readouts per round; one Atto565-conjugated and one Alexa Fluor 647–conjugated), imaging, and TCEP-based reductive cleavage to strip dyes. To reduce total time, each solution change was initiated by priming from source to sample at *∼*0.5 mL min^−1^.

A typical fluidics program per round (excluding priming) was: 3 mL SSC (0.5 mL min^−1^);

1. mL RHB (0.4 mL min^−1^) with round-specific readout probes; a 30 min incubation; 2.5 mL RWB (0.2 mL min^−1^); 2.5 mL 2 *×* SSC (0.2 mL min^−1^); 2 mL IB (0.2 mL min^−1^); then 1 mL IB (0.5 mL min^−1^) and a 15 min incubation prior to imaging. After imaging, 1.3 mL CB (0.13 mL min^−1^) was flowed, followed by a 10 min cleavage incubation, and 3 mL 2 *×* SSC (0.5 mL min^−1^) to rinse.

For cell segmentation and image registration, we first labeled the 20-nt sequence of the poly-dT RNA anchoring probe with a non-cleavable readout conjugated to Alexa Fluor 488 (poly-dT readout), producing a consistent cytoplasmic profile. The general program above was used with an extended overnight hybridization. We also included a set of experiment-unrelated readout probes as blocking oligos to occupy repetitive DNA or other features that might non-specifically bind readouts; these were cleaved of dyes before iterative rounds. We then completed iterative readout staining and imaging as described above.

### Automated multiplexed microscopy

#### Epi-fluorescence microscopy

We constructed a purpose-built widefield microscope for multiplexed RNA-FISH. A 60 *×* NA 1.35 oil-immersion objective (Olympus; UPLXAPO60XO) was mounted to a motorized translation stage (Applied Scientific Instrumentation; LS-50) using a machined mount and XY translator (Thorlabs; LM1XDY). The objective was paired with a 200 mm tube lens (Nikon; MXA20696) to yield an effective magnification of 66 *×*. The system dichroic (Semrock; FF409 / 493 / 573 / 652-Di02) and tube lens were mounted onto a microscope stand (Thorlabs; CFB1500, CSA1002, WFA2002). An industrial CMOS camera (Teledyne FLIR; BFS-U3-200S6M-C) was connected to a right-angle mirror (Thorlabs; KCB2EC, BBE2-E02) and mounted to the op-tical table. A filter wheel (Applied Scientific Instrumentation; FW1000) with three bandpass filters (Semrock; FF01-520/35, FF01-593/40, BLP01-647R) was placed between the tube lens and camera mirror. XY motion used a repurposed sequencing stage and controller (Applied Scientific Instrumentation; MS-2000, LX-4000).

Fluorescence illumination used high-power LEDs (Thorlabs; SOLIS-470C, SOLIS-565C, SOLIS-620D) each driven by a TTL-triggerable controller (Thorlabs; DC20). LEDs were combined using long-pass dichroics (Semrock; FF506-Di03, FF593-Di03). Each LED was spectrally filtered by a bandpass (Semrock; FF01-474/27, FF01-549/15, FF01-635/18) prior to the corresponding dichroic.

Microscope control used a Windows 11 Pro 64-bit workstation (Dell; Precision 3630 Tower) running custom Python software based on Pycro-Manager and Micro-Manager [58, 59]. Hardware control used a programmable DAQ (Austin Blanco Consulting; Triggerscope 4) and system controller (Applied Scientific Instrumentation; LX-4000). LED emission was synchronized to the camera `EXPOSURE OUT` signal to eliminate rolling shutter blur during readout.

#### Buffer exchange system

A peristaltic pump (Gilson; MP1 Single Channel), four 8-way multiplexing valves (Hamilton; MVP, HVXM 8-5), and a 92 µL bubble catcher (Darwin Microfluidics; LVF-KBT-M-A) delivered buffers and readouts. Custom Python code controlled the hardware; programs were stored as text files and integrated into the microscope software. This system was adapted from a Moffitt Lab design.

#### Microscope setup for automated imaging

After poly-dT readout and the smFISH round (readout bits 17 and 18) were hybridized, the sample was loaded onto the XYZ stage. Using the poly-dT–488 readout, the Micro-Manager Stage Explorer plugin generated a large-area overview. A bounding box of the imaging region was defined and tiled with overlap.

Each iterative imaging round comprised readout hybridization, wash, imaging buffer application and imaging, reductive cleavage, and 2 *×* SSC rinse. A representative program was: prime 2 µL RHB over 3 min; run 2.5 µL RHB over 4 min; pause 120 min for hybridization; run 2 µL Buffer D over 3.5 min and then 2 µL Buffer D over 10 min to rinse; run 2 µL imaging buffer over 10 min while priming; push 1 µL Buffer B to advance 1 µL imaging buffer over the sample; pause for imaging. After imaging: run 2 µL CB over 3.5 min to prime, then 1 µL CB over 5 min and pause; then 2 µL Buffer B over 10 min; rinse with an additional 2 µL Wash B over 4 min. This sequence was repeated for each readout round.

### Computing resources

MERFISH simulations were run on a Dell Precision 3660 (Intel i9-13900K, 128 GiB RAM, 8.1 TB storage) running Ubuntu 24.04.3 LTS. All `merfish3d-analysis` results were generated with v0.7.6. Decoding and analysis of MERFISH simulations were performed on Google Colab with an Nvidia A100 GPU. Decoding and analysis of the publicly available dataset and the human olfactory bulb dataset were performed on a home-built server (2 *×* Intel Xeon E5-2650, 1 TB RAM, 2 *×* Nvidia RTX 3090, 100 TB RAID5 storage) running Linux Mint 21 LTS. Local workstation tests were performed using a Lenovo P620 workstation (AMD Threadripper Pro 7945WX, 64 GiB RAM, 4 TB storage, Nvidia RTX 4090) running Ubuntu 24.04.3 LTS.

## Data and Code Availability

`merfish3d-analysis` is maintained on GitHub and version-archived on Zenodo [60]. Raw data for the MERFISH and iterative smFISH simulations are available on Zenodo [61]. The mouse motor cortex bulk RNA sequencing and MERFISH data were downloaded as provided from the data availability section of ref. [45]. The human olfactory bulb codebook, probe sequences, readout sequences are provided as Supplemental File 2. The raw human olfactoy bulb MERFISH data exceed typical file sharing service sizes and are available upon request. A subset of raw human OB MERFISH data is available with the `merfish3d-analysis` documentation.

## Acknowledgements

We acknowledge Dr. Alexis Colloumb for conversations on 3D MERFISH analysis and Dr. Jeffrey Moffitt for providing CAD designs for the automated fluidics reservoir holders plus key advice on performing MERFISH experiments. RK, SJS, MS, PP, SP, DPS acknowledge funding from National Institutes of Health (NIH) RF1MH128867. MA, RK, and BBB acknowledge funding from NIH DP2MH136493. MDS acknowledges support from National Science Foundation Graduate Research Fellowship Program under Grant No. 2233001.

## Author Contributions

DPS created and implemented the software package. MDS, MA, BBB, and DPS tested the software package. MCS, PP, DPS, and SP designed and performed probabilistic MERFISH simulations. DPS designed and implemented the control code for the custom widefield microscopy platform. DPS, SJS, and RK built and validated the custom widefield microscopy platform. RK, SJS, DPS designed and performed MERFISH experiments. MDS, BBB, and DPS performed analysis of iterative smFISH simulations. DPS performed analysis of MER-FISH simulations, public MERFISH data, and MERFISH experiments. RK and DPS wrote the manuscript, with input from all authors.

## Competing Interests

The authors declare no competing interests.

## Notes

https://zenodo.org/records/17274305

https://github.com/QI2lab/merfish3d-analysis

### Competing Interest Statement

The authors have declared no competing interest.

### Summary of Updates

Figure 4 revised and the Methods section updated.

