## Supplemental text for "GPU-accelerated, self-optimizing processing for 3D multiplexed iterative RNA-FISH experiments"

### S1: Probabilistic MERFISH simulations

Here we describe our model to generate realistic synthetic MERFISH data following Eq. 1, replicating the various axial positions, multiple hybridization rounds, and spectral channels seen in a wide-field microscope.

We want to simulate a stack of images with a set of $N$ molecules (of various RNA species), meaning the same set of molecules observed at different focal planes $z_\ell$, with each $\ell \in \{1{:}L\}$ representing the index of a plane in the $z$-stack.

We can index each molecule with $n \in \{1{:}N\}$ across all species and denote the $n$-th particle position as $\vec{x}_n = (x_n, y_n, z_n)$. Each molecule is further assigned a species label $\phi_n \in \{1{:}M\}$, where $M$ is the number of distinct RNA species (or 'barcodes') encoded in the experimental codebook. The codebook defines the activation pattern of each species across $B$ imaging rounds (or bits), with each bit $b \in \{1{:}B\}$ corresponding to a particular spectral channel and hybridization round. Since the mapping between colors, bits, and rounds can vary across experiments, it is convenient to index only by bits, $b$. For simplicity, we will simulate a two-color experiment with alternating colors, where $c = 1$ in all odd bits $b \in 1, 3, 5, \ldots$ and $c = 2$ in all even bits $b \in 2, 4, 6, \ldots$. Therefore, since specifying $b$ will also specify $c$, we will replace color indices $c$ with the bit indices $b$.

Our goal is to simulate MERFISH camera observations, meaning the measured signal of each pixel, each focal plane and each bit in the microscope image with physical accuracy. We index a pixel as $p \in \{1{:}P\}$, and denote by $w^b_{p,\ell}$ the total recorded intensity at pixel $p$ in bit $b$ for the image taken at focal plane $z_\ell$.

Following a standard CMOS camera model, the measured intensity $w^b_{p,\ell}$ is determined by the number of photons incident on the detector element corresponding to pixel $p$, which we denote as $R^b_{p,\ell}$. This photon count is then converted into an analog–digital unit (ADU) through a linear gain factor $\mathscr{G}_p$, combined with a pixel-specific offset $\mu_p$ (the bias level or "dark offset"), and corrupted by additive Gaussian readout noise with variance $v_p$ [62]. Formally,

$$w^b_{p,\ell} \mid R^b_{p,\ell}, \mathscr{G}_p, \mu_p, v_p \sim \text{Normal}\left(\mu_p + \mathscr{G}_p R^b_{p,\ell}, v_p\right). \tag{S1}$$

In order to simulate the number of incident photons per pixel, $R^b_{p,\ell}$, we first decompose it into the sum of background contributions and the contributions of fluorescent probes

$$R^b_{p,\ell} = \xi^b_{p,\ell} + \sum_{n=1}^{N} R^{b,(n)}_{p,\ell}, \tag{S2}$$

where $\xi^b_{p,\ell}$ represents diffuse background photons arriving at pixel $p$ (for example, autofluorescence or out-of-focus light), and $R^{b,(n)}_{p,\ell}$ denotes the number of photons from the RNA molecule $n$ that arrive at pixel $p$.

The background component $\xi^b_{p,\ell}$ in (S2) is introduced to emulate low-spatial-frequency features or uniform autofluorescence sometimes observed in experiments. Thus, we model it as a Poisson random variable with mean proportional to the exposure time $\tau$ and a per-pixel, per-bit background rate $\zeta^b_{p,\ell}$:

$$\xi^b_{p,\ell} \sim \text{Poisson}\left(\tau \zeta^b_{p,\ell}\right). \tag{S3}$$

In contrast, the per-molecule contribution $R^{b,(n)}_{p,\ell}$, each term in the summation in (S2), arises from the fluorescent probes bound to molecule $n$, whose emission is blurred by the optical point spread function (PSF) of the microscope. Formally, we model it as

$$R^{b,(n)}_{p,\ell} \sim \mathrm{Poisson}\Big(\tau \mu^b s^b_n G^b_{p,\ell}(x_n, y_n, z_n)\Big)\,. \tag{S4}$$

We can condense the notation for the sampling of the total number of photons hitting the pixel, $R^b_{p,\ell}$, by substituting (S4) and (S3) into (S2) and using the fact that the sum of independent Poisson random variables is Poisson distributed with mean equal to the sum of means

$$R^b_{p,\ell} \sim \mathrm{Poisson}\left(\tau\left(\zeta^b_{p,\ell} + \sum_{n=1}^{N} \mu^b\, s^b_n\, G^b_{p,\ell}(x_n, y_n, z_n)\right)\right). \tag{S5}$$

Note that each per-molecule contribution (each term in the above sum over $n$) is the product of three factors. First, $\mu^b$ is a predetermined simulation parameter that represents the mean emission rate contributed by a single probe in bit $b$. Next, $s^b_n$ is a random variable that describes the number of probes bound to molecule $n$ in bit $b$, which we will model probabilistically. We assume that each molecule has up to $S^{max}$ available binding sites, and that binding events occur independently across sites with probability $\eta(\phi_n, b)$, which depends on the molecular species $\phi_n$ and the MERFISH codebook. Finally, we have

$$s^b_n \mid \phi_n \sim \mathrm{Binomial}\big(S^{max}, \eta(\phi_n, b)\big)\,, \tag{S6}$$

where $\eta(\phi_n, b)$ takes values $\eta_{\mathrm{on}}$ or $\eta_{\mathrm{off}}$ depending on whether bit $b$ is active or inactive for species $\phi_n$. Importantly, the binding probability $\eta(\phi_n, b)$ is the only place where the MERFISH codebook enters our model. If species $\phi_n$ has barcode $\mathscr{C}_{\phi_n} = [\mathscr{C}_{\phi_n,1}, \ldots, \mathscr{C}_{\phi_n,B}]$ with $\mathscr{C}_{\phi_n,b} \in \{0, 1\}$, then

$$\eta(\phi_n, b) = \begin{cases} \eta_{\mathrm{off}}, & \text{if} \quad \mathscr{C}_{\phi_n,b} = 0, \\ \eta_{\mathrm{on}}, & \text{if} \quad \mathscr{C}_{\phi_n,b} = 1, \end{cases} \tag{S7}$$

where $\eta_{\mathrm{off}}$ and $\eta_{\mathrm{on}}$ denote the probability of probe binding for inactive and active codebook entries, respectively. In an ideal experiment, $\eta_{\mathrm{off}} = 0$ and $\eta_{\mathrm{on}} = 1$, corresponding to perfectly specific probes with full occupancy of target sites. However, to better capture realistic imaging conditions, we instead use $\eta_{\mathrm{off}} = 0.01$ and $\eta_{\mathrm{on}} = 0.5$, reflecting a small amount of off-target binding and variable on-target binding efficiency. We further set $S^{max} = 90$ binding sites per RNA molecule, consistent with previous probe designs [10]. Then, to account for off-target binding, we model fluorescent spots that do not correspond to true RNA molecules as an additional "pseudo-species" $\phi_n = M + 1$. These anomalous features are assigned the pseudo-barcode $\mathscr{C}_{M+1} = [2, 2, \ldots, 2]$, and their binding probability is defined as

$$\eta(\phi_n, b) = \begin{cases} \eta_{\mathrm{off}}, & \text{if} \quad \mathscr{C}_{\phi_n,b} = 0, \\ \eta_{\mathrm{on}}, & \text{if} \quad \mathscr{C}_{\phi_n,b} = 1, \\ \eta_{\mathrm{anomalous}}, & \text{if} \quad \mathscr{C}_{\phi_n,b} = 2, \end{cases} \tag{S8}$$

where $\eta_{\mathrm{anomalous}}$ governs the frequency of anomalous binding events.

Finally, $G^b_{p,\ell}(x_n, y_n, z_n)$ in (S4) quantifies how much of the emission from molecule $n$ contributes to pixel $p$ when the image is taken at focal plane $z_\ell$. This factor is obtained by integrating the object-space PSF $g^b$ of the microscope over the finite area of pixel $p$ (centered at $(x_p, y_p)$ with width $\delta_{\text{pix}}$):

$$G^b_{p,\ell}\left(x_n, y_n, z_n\right) = \int_{x^p - \frac{\delta_{\text{pix}}}{2}}^{x^p + \frac{\delta_{\text{pix}}}{2}} \int_{y^p - \frac{\delta_{\text{pix}}}{2}}^{y^p + \frac{\delta_{\text{pix}}}{2}} g^b\left(x_n, y_n, z_n; x, y, z_\ell\right)\, dx\, dy. \tag{S9}$$

This formulation captures the spatial blurring introduced by the optics and ensures that molecular emissions are distributed across neighboring pixels according to the microscope's resolution.

In summary, given the molecular positions $\{\vec{x}_n\}_{n=1}^N$, the species assignments $\{\phi_n\}_{n=1}^N$, the focal plane heights $\{z_\ell\}_{\ell=1}^L$, the binding probabilities $\eta(\phi_n, b)$ specified by the codebook, the background intensity field $\{\zeta^b_{p,\ell}\}$, and the camera parameters $(\tau, \mu^b, \mathscr{G}_p, \mu_p, v_p)$, the image formation process can be summarized by the following set of sampling equations:

$$\eta(\phi_n, b) = \begin{cases} \eta_{\text{off}}, & \text{if} \quad \mathscr{C}_{\phi_n, b} = 0, \\ \eta_{\text{on}}, & \text{if} \quad \mathscr{C}_{\phi_n, b} = 1, \\ \eta_{\text{anomalous}}, & \text{if} \quad \mathscr{C}_{\phi_n, b} = 2, \end{cases} \tag{S10a}$$

$$s^b_n \mid \phi_n \sim \text{Binomial}\left(S^{max}, \eta(\phi_n, b)\right), \tag{S10b}$$

$$R^b_{p,\ell} \sim \text{Poisson}\left(\tau\left(\zeta^b_{p,\ell} + \sum_{n=1}^{N} \mu^b s^b_n G^b_{p,\ell}(x_n, y_n, z_n)\right)\right), \tag{S10c}$$

$$w^b_{p,\ell} \mid R^b_{p,\ell}, \mathscr{G}_p, \mu_p, v_p \sim \text{Normal}\left(\mu_p + \mathscr{G}_p R^b_{p,\ell},\ v_p\right). \tag{S10d}$$

This set of sampling equations completes our image simulation.

### S2: Detailed decoding of individual tiles from mouse motor cortex MERFISH data

Re-analyzing the publicly available mouse motor cortex data revealed a key challenge for analysis replication. First, the actual raw data is not available, meaning we had to begin with a partially processed dataset with an unknown interpolation applied. Second, the complete set of parameters used for decoding was not stored with the data, leading to uncertainty in how to appropriately re-analyze. Finally, we found that some of the recorded stage transformations were incorrect, and we had to manually determine the relative tile placement before proceeding with the analysis of the entire dataset.

To re-analyze the publicly available data, we made the following assumptions:

1. Based on searching the `merlin` codebase for default values, we used a lower pixel magnitude of 0.9 to match their analysis.

2. We had to manually optimize both the initial $(x, y)$ positions and stage orientations, beginning from the stage positions made available with the data. Without these optimizations, it was not possible to globally registered the data.

3. The README associated with the publicly available datset states "Note: For the 650 nm channels, a significant number of spots observed the first z-plane (i.e. at the coverslip surface) correspond to non-specific binding of the 650 nm dye to the coverslip surface, and the vast majority of these non-specific binding spots are decoded as invalid barcodes in the decoding process and are not used for subsequent analysis." [45]

4. The raw data contains z planes at the coordinates

$$[0, 1.5, 3, 4.5, 6, 7.5]$$

Even with these assumptions and excluded data at the coverslip, we could not recover the same results using `merlin` and `merfish3d-analysis`. After further examining the publicly available data, we found a number of pixels outside cell bodies where `merlin` placed a decoded RNA molecule. Upon closer inspection, the data contained minimal evidence in either the magnitude values or distance values that a labeled RNA was present in that Z plane, or the immediately adjacent Z planes. In contrast, `merfish3d-analysis` does not place at an RNA molecule at these pixels, contributing to the observed discrepancy in the number of RNA localized outside of cells between the two approaches (Figures S1, S2). While we were unable to independently perform the analysis using `merlin` to confirm if these RNA localizations are spurious and should have been removed, this subtlety in RNA localization highlights the complexities associated with decoding and subsequently sharing such large datasets.

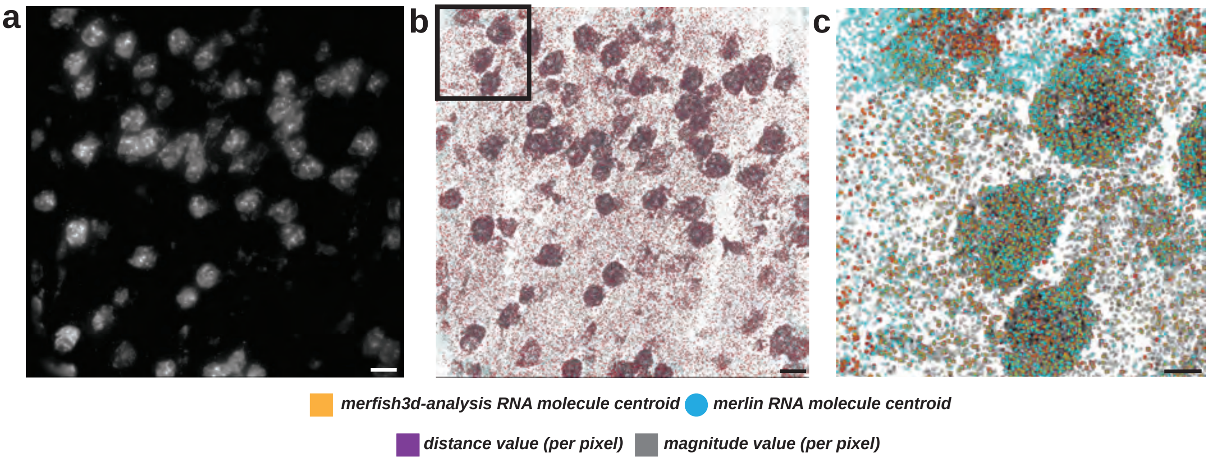


Figure S1. Decoding results for tile 40 Decoding results for tile 40 in the publicly available dataset. a) Maximum z projection of deconvolved polyDT stain. b) Maximum z projection of `merlin` (cyan circles) and `merfish3d-analysis` (orange squares) overlaid on pixel distance values (magenta) and pixel magnitude values (inverted grayscale) for the entire tile. Inset shows area displayed in (c). c) Decoding results for maximum z projection of highlighted area in (b). Scale bars (a-b) 15 µm, (c-e) 5 µm.

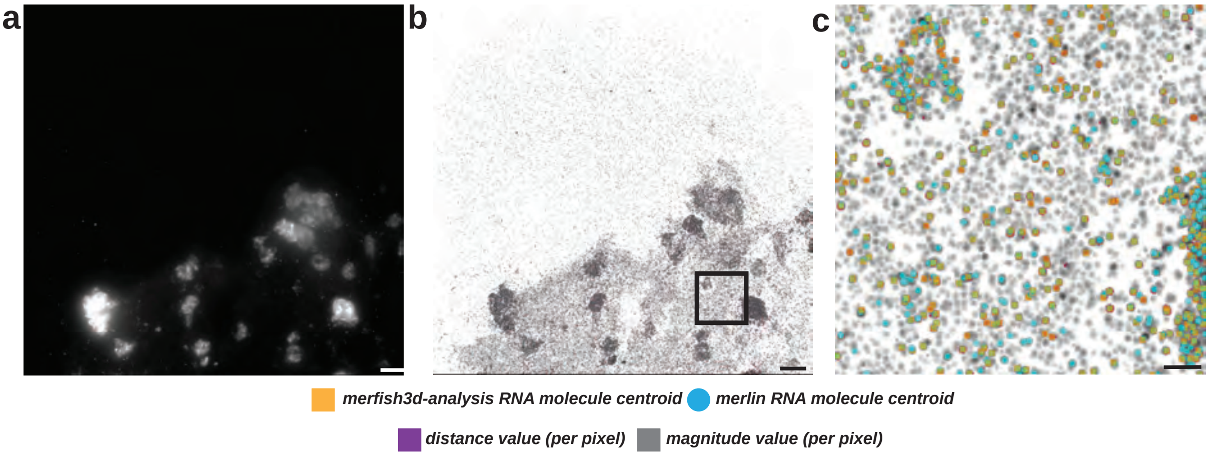


Figure S2. Decoding results for tile 53 Decoding results for tile 53 in the publicly available dataset. a) Maximum z projection of deconvolved polyDT stain. b) Maximum z projection of `merlin` (cyan circles) and `merfish3d-analysis` (orange squares) overlaid on pixel distance values (magenta) and pixel magnitude values (inverted grayscale) for the entire tile. Inset shows area displayed in (c). c) Decoding results for maximum z projection of highlighted area in (b). Scale bars (a-b) 15 µm, (c-e) 5 µm.

### S3: Iterative smFISH

To explore a potential extension of the `merfish3d-analysis` framework to non-error correcting codebooks, we simulated an iterative smFISH experiment, using the same simulation framework as the MERFISH simulation, instead using a 10-bit iterative smFISH codebook encoding codewords for 10 species with 0 blank controls and simulated two spatial distribution of ground truth RNA molecules locations to test the effect of crowding. In the "uniform" condition, we randomly distributed RNA molecules throughout the simulation volume. In the "cell" condition, we primarily randomly distributed RNA molecules inside 3D cylinders with the typical volume of a cell ($\sim$ 10 µm diameter).

We then took these two ground truth simulations and generated synthetic microscopy data at three different axial spacings, 0.315 µm, 1.0 µm and 1.5 µm. For each imaging round, we simulated diffraction limited fiducial markers and two bits of the codebook with a detailed, probabilistic model that accounted for both biochemical and measurement noise parameters (Methods, Supplement Section 1). The iterative smFISH simulations parameters are summarized in Table S2.

| | Uniform | Cells |
|---|---|---|
| Number of RNA species | 10 | 10 |
| Total RNA molecules | 3333 | 1500 in cells; 200 outside |
| Number of molecules per RNA species | $\sim$333 | $\sim$150 |
| Fluorescent spot density (molecule per $\mu m^2$) | $\sim 5 \cdot 10^{-2}$ | $\sim 4 \cdot 10^{-2}$ in cells |

Table S2. smFISH simulation details

For all simulations, we ran the standard `merfish3d-analysis` pipeline as above, modifying the distance threshold to1.0 and allowing for pixels with magnitude thresholds from 0.75 to 1.5. These values were determined through empirical optimization of decoding parameters using the known ground truth RNA locations and identity. The decoding results are summarized in Table Table S2 and Figures Figures S3, S4. For both spatial simulation types, we found reasonable recovery of ground truth RNA molecule locations and identity for 0.315 µm and 1.0 µm axial spacings and a loss of $\sim$ 45 % of RNA molecules for 1.5 µm axial spacing.

| Simulation Type | Axial Spacing | Precision | Recall | F1 score | Figure |
|---|---|---|---|---|---|
| Uniform smFISH | 0.315 µm | 0.88 | 0.86 | 0.86 | S3 |
| Uniform smFISH | 1.0 µm | 0.69 | 0.86 | 0.65 | S3 |
| Uniform smFISH | 1.5 µm | 0.61 | 0.54 | 0.57 | S3 |
| Cells smFISH | 0.315 µm | 0.85 | 0.65 | 0.74 | S4 |
| Cells smFISH | 1.0 µm | 0.69 | 0.65 | 0.66 | S4 |
| Cells smFISH | 1.5 µm | 0.59 | 0.52 | 0.55 | S4 |

Table S2. Summary statistics of decoding results for iterative smFISH simulations Decoding results for iterative smFISH simulation of randomly distributed RNA molecules (Uniform smFISH) and iterative smFISH simulation of RNA molecules within cells (Cells smFISH) at three different z spacings 0.315 µm, 1.0 µm and 1.5 µm using `merfish3d-analysis` with the default parameters, except for distance threshold (1.0) and magnitude threshold (1.5). All values rounded to two significant digits.

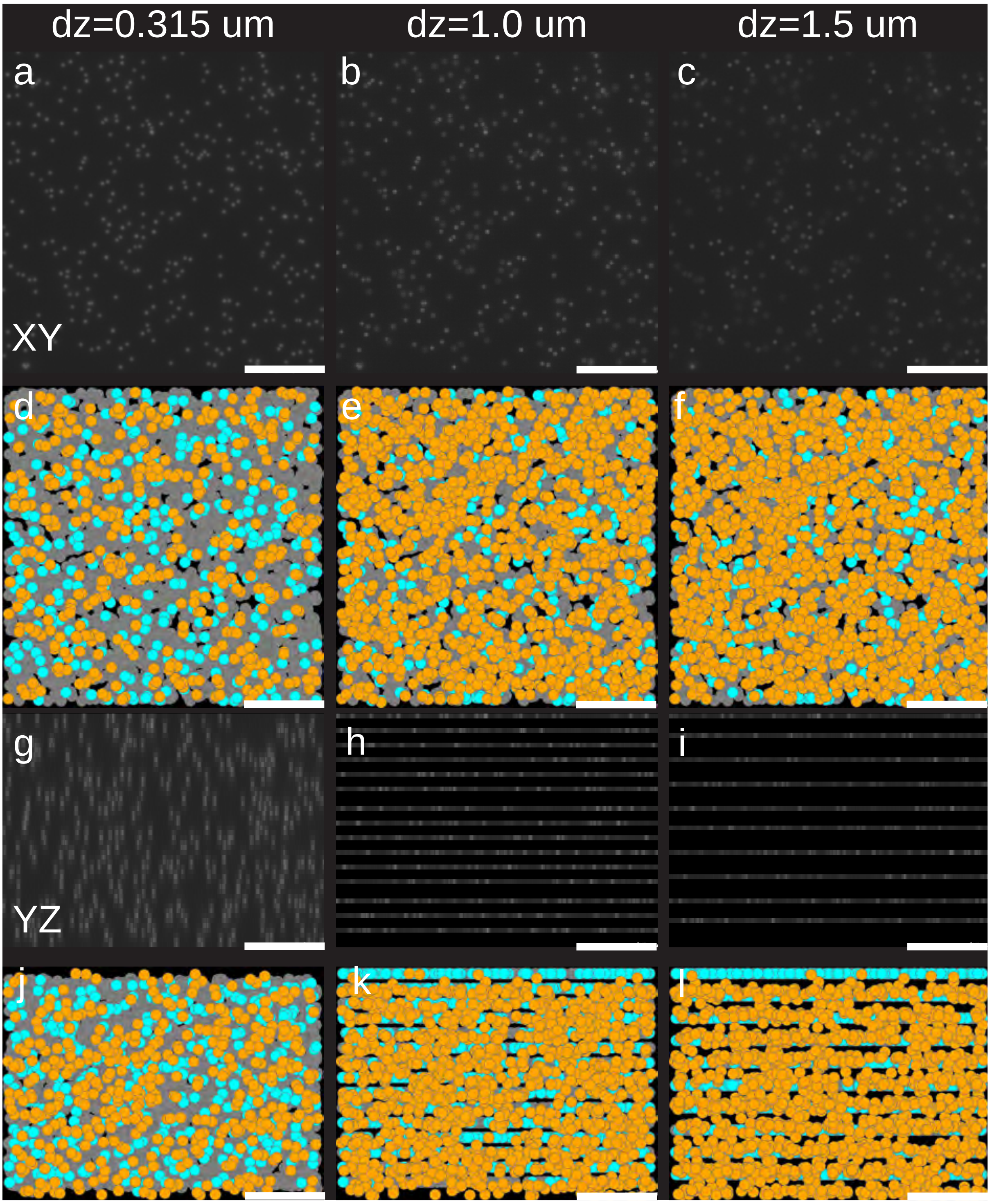


Figure S3. Uniform distributed smFISH simulation results Maximum Z projections (XY view) of one bit for a) dz =0.315 µm, b) dz =1.0 µm, and c) dz =1.5 µm axial sampling. Maximum Z projection(XY view) of recovered RNA true positives (gray), false positives (cyan), and false negatives (orange) for d) dz =0.315 µm, e) dz =1.0 µm, and f) dz =1.5 µm axial sampling. Maximum X projections (YZ view) of one bit for g) dz =0.315 µm, h) dz =1.0 µm, and i) dz =1.5 µm axial sampling. Maximum X projections (YZ view) of recovered RNA true positives (gray), false positives (cyan), and false negatives (orange) for j) dz =0.315 µm, k) dz =1.0 µm, and l) dz =1.5 µm axial steps. All scale bars 5 µm.

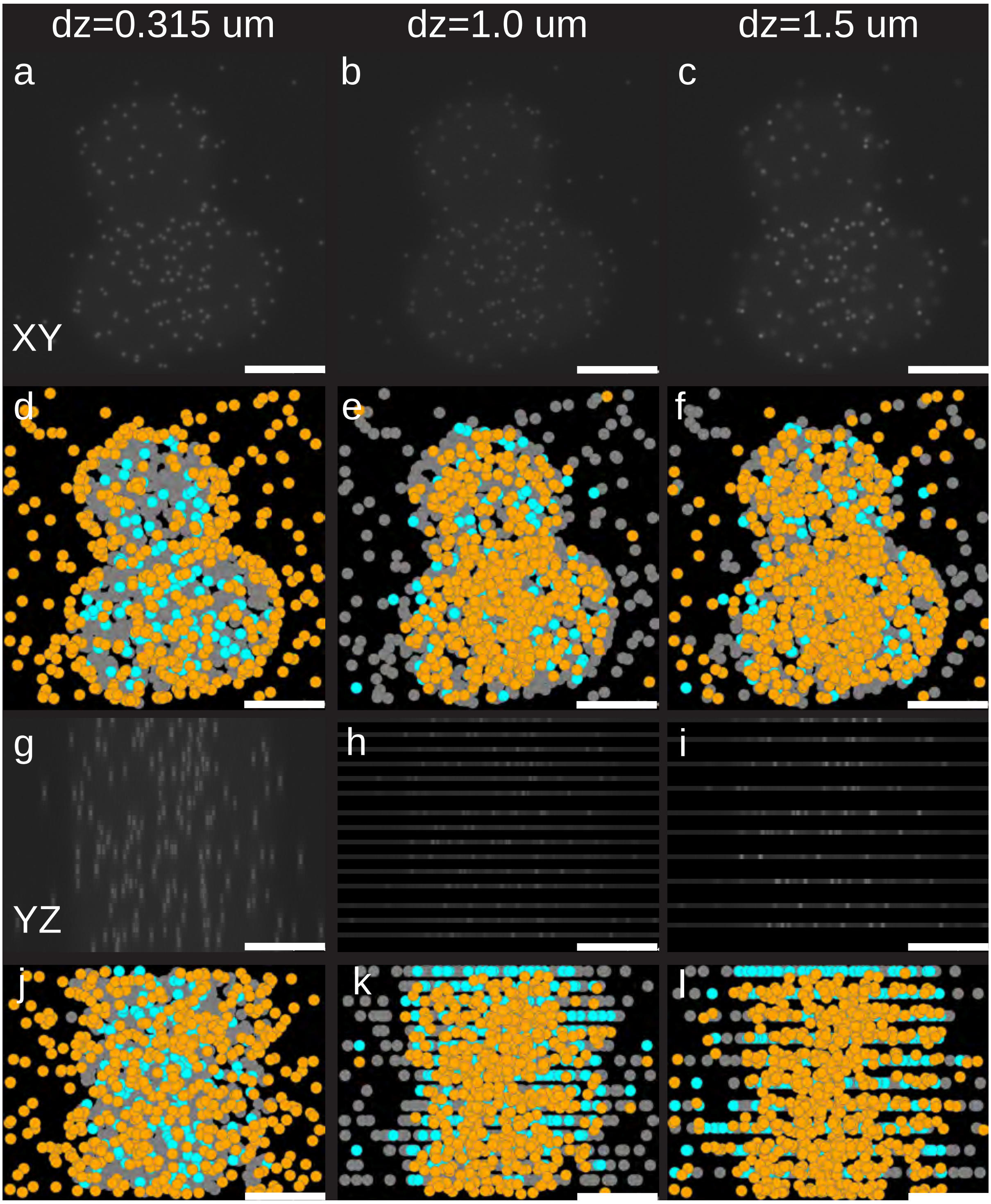


Figure S4. Cell distributed smFISH simulation results Maximum Z projections (XY view) of one bit for a) dz =0.315 µm, b) dz =1.0 µm, and c) dz =1.5 µm axial sampling. Maximum Z projection(XY view) of recovered RNA true positives (gray), false positives (cyan), and false negatives (orange) for d) dz =0.315 µm, e) dz =1.0 µm, and f) dz =1.5 µm axial sampling. Maximum X projections (YZ view) of one bit for g) dz =0.315 µm, h) dz =1.0 µm, and i) dz =1.5 µm axial sampling. Maximum X projections (YZ view) of recovered RNA true positives (gray), false positives (cyan), and false negatives (orange) for j) dz =0.315 µm, k) dz =1.0 µm, and l) dz =1.5 µm axial steps. All scale bars 5 µm.
